# Genome-wide methylation data mirror ancestry information

**DOI:** 10.1101/066340

**Authors:** Elior Rahmani, Liat Shenhav, Regev Schweiger, Paul Yousefi, Karen Huen, Brenda Eskenazi, Celeste Eng, Scott Huntsman, Donglei Hu, Joshua Galanter, Sam Oh, Melanie Waldenberger, Konstantin Strauch, Harald Grallert, Thomas Meitinger, Christian Gieger, Nina Holland, Esteban Burchard, Noah Zaitlen, Eran Halperin

## Abstract

Genetic data are known to harbor information about human demographics, and genotyping data are commonly used for capturing ancestry information by leveraging genome-wide differences between populations. In contrast, it is not clear to what extent population structure is captured by whole-genome DNA methylation data. We demonstrate, using three large cohort 450K methylation array data sets, that ancestry information signal is mirrored in genome-wide DNA methylation data, and that it can be further isolated more effectively by leveraging the correlation structure of CpGs with cis-located SNPs. Based on these insights, we propose a method, EPISTRUCTURE, for the inference of ancestry from methylation data, without the need for genotype data. EPISTRUCTURE can be used to infer ancestry information of individuals based on their methylation data in the absence of corresponding genetic data. Although genetic data are often collected in epigenetic studies of large cohorts, these are typically not made publicly available, making the application of EPISTRUCTURE especially useful for anyone working on public data. Implementation of EPISTRUCTURE is available in GLINT, our recently released toolset for DNA methylation analysis at: http://glint-epigenetics.readthedocs.io.

## Background

The relation between ancestry and genetic variation has been repeatedly established over the last decade [1,2]. Several methods now provide accurate estimates of ancestry information by leveraging genome-wide systematic difference in allele frequencies between subpopulations, commonly referred to as population structure [3–7]. These methods often apply Principal Component Analysis (PCA) or variants of PCA.

Inferring population structure across individuals provides a powerful source of information for various fields, including genetic epidemiology, pharmacogenomics and anthropology. For instance, in genetic and molecular epidemiology, in which identifying genetic associations with disease or exposure is of primary interest, it is essential to have ancestry information in order to distinguish effects of demographic processes from biological or environmental effects. Specifically, the importance of controlling for population structure in genome-wide association studies (GWAS) is now well appreciated. Unless appropriately accounted for, population structure in GWAS can lead to numerous spurious associations and might obscure true signals [4,8].

Emerging epigenome-wide association studies (EWAS) revealed thousands of CpG methylation sites correlated with genetics and with ancestry [9–20]. Not surprisingly, due to the genetic signal present in many CpGs, several studies have shown that the first several principal components (PCs) of methylation data can capture population structure in cohorts composed of European and African individuals [15,21]. However, unlike the case of genotyping data in which global ancestry information is robustly reflected by the top PCs, the first several PCs of methylation data were also shown to capture other factors in different scenarios, mainly cell type composition in case of data collected from heterogeneous tissues [22,23], but also other factors, including technical variables, age and sex [15,21]. Moreover, it is now appreciated that collecting methylation using probes with polymorphic CpGs is affected by hybridization sensitivity and does not necessarily reflect methylation variability but rather genetic variability [24]. Therefore, it is not clear to what extent global whole-genome DNA methylation states are affected by population structure when these artifacts are removed.

We hereby introduce EPISTRUCTURE, a method for capturing ancestry information from DNA methylation data. EPISTRUCTURE is based on the observation that PCA computed from a set of methylation CpG sites that are highly correlated with SNPs efficiently captures population structure. Thus, we use a large reference data set that includes both genotypes and methylation in order to find correlations of CpGs with cis-located SNPs, and to generate a reference list of genetically-informative CpGs. Then, given new methylation data we compute the PCs of the methylation levels from the same sites included in the reference list. We validate the robustness of this method by assessing the correlation between the methylation-inferred ancestry and the genetically inferred ancestry on two additional large methylation data sets.

In order to shed light on the relation between genetic ancestry and methylation-based ancestry, we further explore the unsupervised detection of ancestry from methylation data. We show that genome-wide methylation mirrors ancestry information in admixed populations after properly adjusting for known variability in genome-wide methylation, and after properly removing technical artifacts, particularly probes that include SNPs that may confound the results. Thus, unlike previous studies that were potentially confounded by these artifacts, here we show that ancestry is indeed robustly mirrored by methylation data as one of the main variance components in the data.

EPISTRUCTURE can be used to infer ancestry information from methylation data in the absence of genetic data. Although genetic data are often collected in epigenetic studies of large cohorts, these are typically not made publicly available, making the application of EPISTRUCTURE especially useful for anyone working on public data.

## Results

### Inferring ancestry from methylation data using EPISTRUCTURE

Ancestry information signal in methylation is mostly expected to exist due to the large number of correlations between methylation sites and genetics [9–20]. We developed EPISTRUCTURE, a method for the inference of ancestry from methylation data, which relies on reference data in which both genotype and methylation data are available. Briefly, EPISTRUCTURE selects a set of CpGs that are highly correlated with genotype information in the reference data, and then, given new methylation data, performs principal component analysis on these sites while taking into account the cell type composition effects. More specifically, we use the KORA cohort of European adults (n=1,799), for which both whole-blood methylation and genotyping data are available [25] (see Methods). We fitted a regularized linear regression model for each CpG from SNPs in cis, and evaluated it based on a cross-validated linear correlation (see Methods). Since the vast majority of reported CpG-SNP associations are between CpGs and cis-located SNPs [9–11], we only considered cis-located SNPs in capturing the genetic component of each CpG. We observed that for most CpGs only a small fraction of the variance can be explained by cis-SNPs (*R*^2^ < 0.1 for 92.9% of the CpGs tested; Supplementary Figure S1), thus motivating the use of only a relatively small subset of the CpGs for inferring ancestry information. Considering only sites that most of their variance can be explained by cis-SNPs (*R*^2^ > 0.5) resulted in a reference list of 4,913 genetically-informative CpGs (see Methods). We note that polymorphic CpGs were not excluded from the KORA data set before learning the reference of informative CpGs, therefore polymorphic CpGs that can be well explained by cis-SNPs (*R*^2^ > 0.5) were also included in the reference list. In most cases, polymorphic CpGs should be excluded before any data analysis, however, in our case, EPISTRUCTURE leverages the true genetic signal underlying in the probes of these CpGs. We later demonstrate the difference in performance when excluding these probes.

In order to test the performance of EPISTRUCTURE we applied it on the GALA II data set (*n* = 479), a pediatric Latino population study with Mexican (MX) and Puerto-Rican (PR) individuals [26], for which both genotypes and 450K methylation array data (whole-blood) were available (see Methods). First, we computed the largest (first) two PCs of the genotypes (genotype-based PCs), known to capture population structure [4]. We observed that the first PC of EPISTRUCTURE captured the top genotype-based PC well (*R*^2^ = 0.82), as compared to the first PC of the methylation data (methylation-based PC; *R*^2^ = 0.01) and as compared to the methylation-based PC computed only from CpGs residing in close proximity to nearby SNPs (*R*^2^ = 0.01), as was suggested in a recent study for capturing ancestry information in methylation data [21]. Further, we observed that EPISTRUCTURE provides substantially improved correlation with the first two genotype-based PCs as compared with the alternatives (Figure 1).

**Figure 1:**
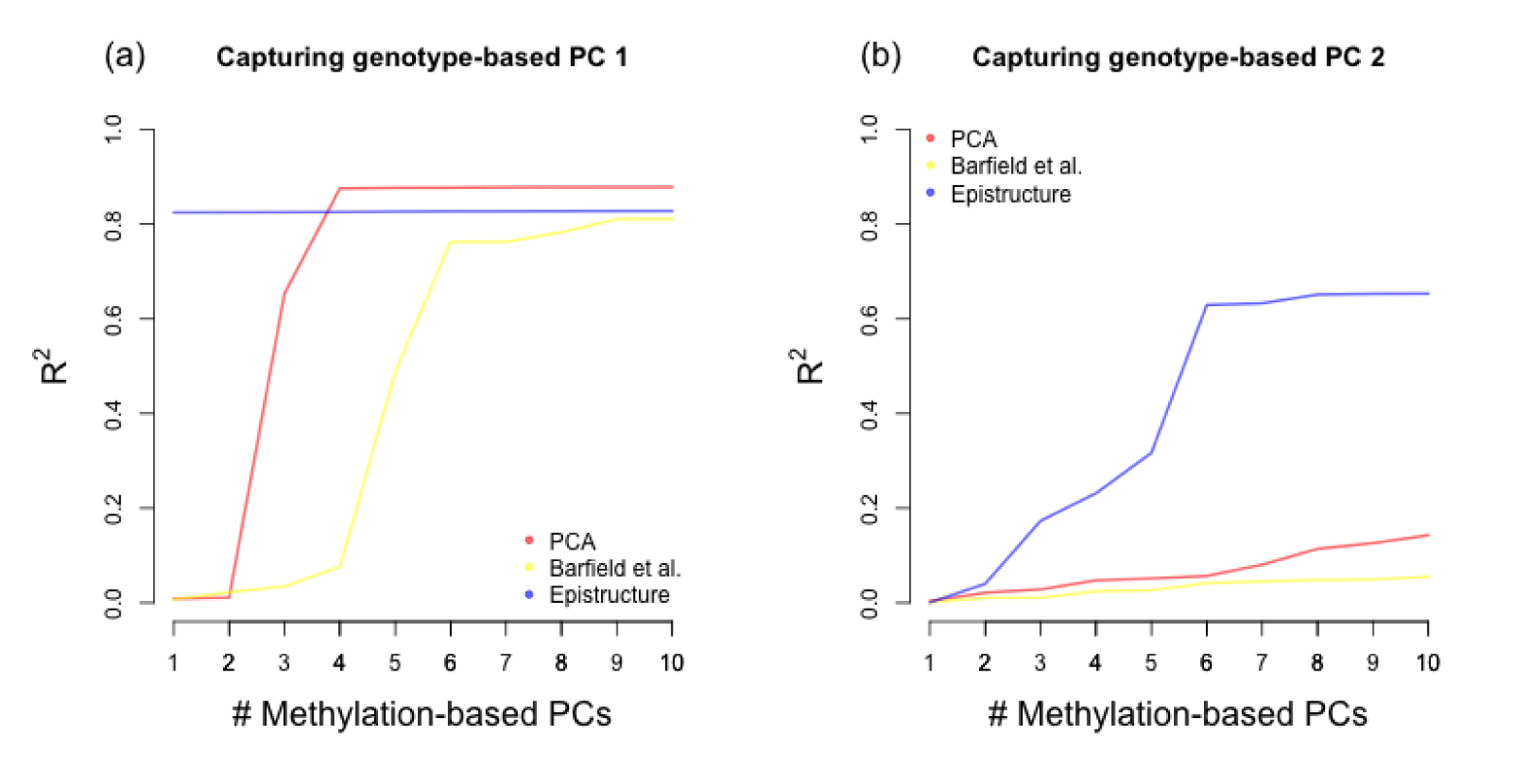
The fraction of variance explained in the first two genotype-based PCs of the GALA II data using several methods. Presented are linear predictors using increasing number of EPISTRUCTURE PCs (in blue), methylation-based PCs (in red) and methylation-based PCs after feature selection based on a previous study [21] (in yellow) for capturing (a) the first genotype-based PC and (b) the second genotype-based PC.

Next, as an alternative measure of population structure, we used the ADMIXTURE software [5] to estimate, for each individual, ancestry proportions of the three ancestries known to compose the MX and PR populations: European (EU), Native-American (NA) and African (AF). In this case, the top two principal components of EPISTRUCTURE capture very well both the Native American ancestry and the African Ancestry (*R*^2^ = 0.81 and *R*^2^ = 0.56 respectively), while the European ancestry was captured to a lesser extent (*R*^2^ = 0.32, see Supplementary Figure S2).

We further tested whether ancestry information can be captured using EPISTRUCTURE in case there is a weaker population structure in the data. We observed that the first two PCs of EPISTRUCTURE could capture ancestry information well in both subpopulations of the GALA II data (*R*^2^ = 0.33 in the PR group and *R*^2^ = 0.76 in the MX group; Supplementary Figure S3). These results suggest that EPISTRUCTURE can be used as an easy and efficient method for capturing ancestry information in methylation, even in data sets with relatively modest population structure.

### Unsupervised ancestry inference from methylation data

EPISTRUCTURE is a supervised approach since it uses a reference data set in which both methylation and genotype data are available. In order to shed light on the extent to which ancestry is reflected by methylation data, we also explored unsupervised approaches for the inference of ancestry from methylation data. Consistent with a previous study of individuals from the same population [27], the first two genotype-based PCs of the GALA II data clustered the samples into two groups, generally corresponding to MX and PR subpopulations (Figure 2a). Since PCA has been shown to mirror ancestry very accurately in the case of genetic data [1], we first computed the top two methylation-based PCs while accounting for known technical factors as well as for age and sex, which are known to affect methylation genome-wide [28–30]. Considering the population structure characterized by the first two genotype-based PCs as the “gold standard”, the first two methylation-based PCs could not sufficiently capture the population structure in the data (Figure 2b).

**Figure 2:**
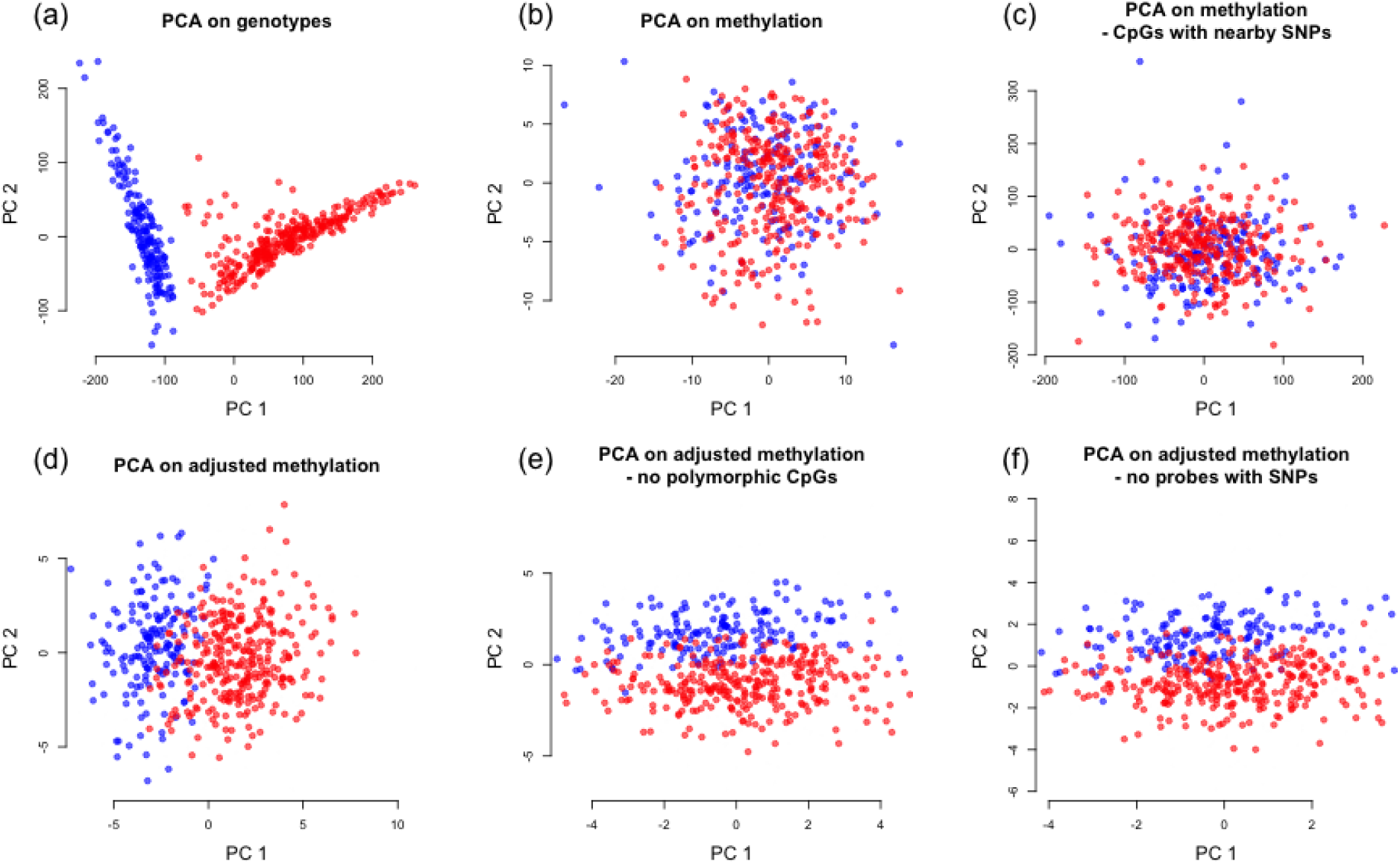
Capturing population structure in the GALA II data using an unsupervised approach. (a) The first two PCs of the genotypes, considered as the gold standard, separate the samples into two subpopulations: Puerto-Ricans (in blue) and Mexicans (in red). (b) The first two PCs of the methylation levels (methylation PCs) cannot reconstruct the separation found with the genotype data. (c) Recalculating the first two PCs after applying a feature selection based on proximity of CpGs to nearby SNPs as was proposed by Barfield et al. [21] (d) The first two PCs of the methylation after adjusting the data for cell type composition (adjusted methylation PCs) can reconstruct most of the separation found in the genotypes. (e) Using adjusted methylation PCs after excluding the 70,889 polymorphic sites from data. (f) Using adjusted methylation PCs after excluding the 167,738 probes containing at least one common SNP.

We then applied a few more sophisticated procedures, as follows. First, as before, we applied a feature selection step prior to calculating the methylation-based PCs according to a recent study, that suggested to consider only CpGs residing in close proximity to nearby SNPs in order to capture ancestry information in the first few PCs of the data [21]. We found that this procedure did not sufficiently reflect population structure in methylation data (Figure 2c). Next, since the first several PCs in methylation data coming from heterogeneous source such as blood are known to be dominated by cell type composition [22, 23, 31], we adjusted the data for cell type composition using the ReFACTor software [23] and recalculated the first two PCs. This approach effectively reconstructed most of the separation determined by the genotype-based PCs (Figure 2d).

These results show that 450K-probed methylation data indeed reflect genotype data well. Specifically, after accounting properly for known confounders, the top methylation-based PCs capture the genotype-based PCs. However, these results can potentially be driven by artifacts. Specifically, it is now acknowledged that many probes in the 450K methylation array contain single nucleotide polymorphisms (SNPs) in their target CpGs. Such polymorphic CpGs were shown to bias measured methylation levels as a function of the individual’s genotypes, apparently due to changes in probe binding specificity [24]. Thus, our results so far might be biased by these probes. To address this possibility, we recalculated the first two methylation-based PCs after excluding 70,889 CpGs that are known to be polymorphic (see Methods). We found that the new methylation-based PCs could still capture well the first genotype-based PC (*R*^2^ = 0.77 as opposed to *R*^2^ = 0.83 before removing the polymorphic CpGs), accounting for the separation found using the first two genotype-based PCs (Figure 2e). In addition, we performed a more conservative analysis by repeating the last step after further excluding 167,738 probes containing at least one common SNP anywhere in the probe (i.e. not only in the target CpG; see Methods). We found that in this case as well the reconstruction using the top two methylation-based PCs provided almost the same separation determined by the genotype-based PCs (Figure 2f; *R*^2^ = 0.70 with the first genotype-based PC).

We note that repeating the last two experiments while accounting for estimated cell proportions computed using a commonly applied reference-based method [32] as an alternative approach for correction of cell composition effects in methylation could not achieve the same results (*R*^2^ = 0.23 and *R*^2^ = 0.14 in the experiments without the polymorphic sites and in the experiment removing all probes with common SNPs, respectively). This can be explained by the additional cell type composition signal captured by ReFACTor but not by the reference-based approach, as was previously demonstrated on the GALA II data [23]. Substantial difference in performance is especially expected in cases where the reference methylation data used by the reference-based method do not represent the target population well [23,33]. Removing only part of the cell type composition signal from the data results in PCs that are likely to be still dominated by tissue composition information rather than by population structure. Alternatively, it may be the case that ReFACTor also removed another sparse confounder, in addition to the cell type composition signal.

We also compared the different approaches using the ancestry estimates of the ADMIXTURE software [5]. The results were consistent with our previous experiment – while the first two methylation-based PCs could not capture the ancestry estimates (*R*^2^ = 0.02 with the EU fraction, *R*^2^ = 0.01 with NA and *R*^2^ = 0.02 with AF), we found the first two methylation-based PCs after adjusting for cell composition to capture the ancestry estimates well, even after excluding from the data all probes containing common SNPs (*R*^2^ = 0.28 with the EU fraction, *R*^2^ = 0.69 with NA and *R*^2^ = 0.47 with AF; Supplementary Figure S4).

We further tested whether ancestry information can be captured in the same manner when applied to each of the two subpopulations in the data (MX and PR) separately. We found the methylation-based PCs to capture well only the first genotype-based PC of the Mexican group when not excluding probes containing common SNPs (*R*^2^ = 0.08 for the PR cluster and *R*^2^ = 0.74 for the MX cluster). After excluding the 167,738 probes containing at least one common SNP from the data, the methylation-based PCs could not capture a substantial fraction of the first genotype-based PC in either clusters (*R*^2^ = 0.05 for the PR cluster and *R*^2^ = 0.05 also for the MX cluster). Thus, we conclude that under weak population structure the current unsupervised approach do not mirror ancestry well. However, as we demonstrated earlier, the supervised approach of EPISTRUCTURE, using only a relatively small subset of highly informative CpGs (including highly informative polymorphic CpGs), performed well in this case.

### Validation using the CHAMACOS study data

We further validated the effectiveness of EPISTRUCTURE and the unsupervised approaches using data from the primarily Mexican-American CHAMACOS cohort [34,35]. We used whole-blood methylation levels from nine years old participants (n=227) for which we had 106 ancestry informative markers (AIMs) [36], previously shown to approximate ancestry information well in another Hispanic admixed population [37].

We computed the first two PCs of the available AIMs (genotype-based PCs) in order to capture the ancestry information of the samples. Since the CHAMACOS cohort primarily consists of Mexican-American individuals, we observed no separation into distinct subpopulations in the first several genotype-based PCs. We then computed the first two methylation-based PCs, before and after adjusting the data for cell composition. In addition, we computed the first two EPISTRUCTURE PCs of the data, and measured how much of the variance of the first genotype-based PC can be explained by each of the approaches. As shown in Figure 3, the first two methylation-based PCs could capture only a small portion of the first genotype-based PC (*R*^2^ = 0.04 before adjusting for cell composition and *R*^2^ = 0.16 after adjusting for cell composition), as opposed with the first two EPISTRUCTURE PCs which could capture the first genotype-based PC substantially better (*R*^2^ = 0.38). As in the GALA II data, applying feature selection based on proximity of CpGs to SNPs [21] could capture only a small portion of the ancestry information (*R*^2^ = 0.05).

**Figure 3:**
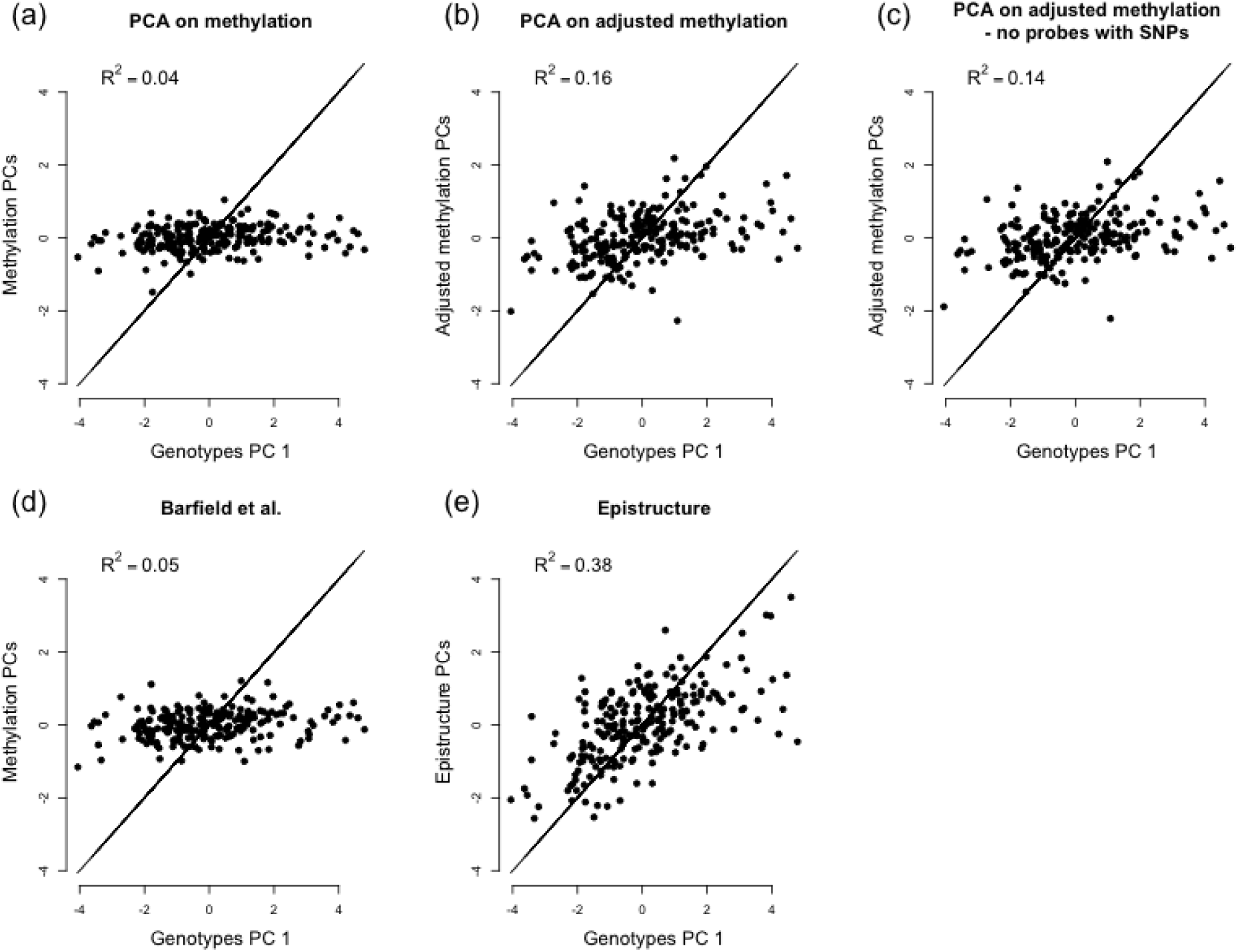
Capturing population structure in the CHAMACOS data. Presented are linear predictors of the first genotype-based PC using (a) the first two methylation PCs of the data, (b) the first two PCs after adjusting the data for cell type composition (adjusted methylation PCs), (c) the first two adjusted methylation PCs after excluding 167,738 probes containing SNPs from the data, (d) the first two PCs calculated based on a previously suggested feature selection [21] and (e) the using the first two EPISTRUCTURE PCs.

As before, we used the ADMIXTURE software [5] as an alternative measure of population structure. For each individual we estimated the ancestry proportions of the three ancestries known to compose Mexican individuals: European (EU), Native-American (NA) and African (AF). The first two EPISTRUCTURE PCs were found to explain a large portion of the EU and NA fraction estimates (*R*^2^ = 0.46 for EU and *R*^2^ = 0.6 for NA ancestry), as opposed with the first two methylation-based PCs (*R*^2^ = 0.11 for EU and *R*^2^ = 0.14 for NA ancestry, after adjusting for cell type composition; Supplementary Figure S5). The estimates of African proportions, however, were not captured well by either approach. This result was expected due to the low average proportion of African ancestry in Mexican samples (less than 10%) [38]. All the results are summarized in Supplementary Table S1.

### Implications for the EPIC (850K) array

The recently introduced EPIC array by Illumina, which allows to probe a set of approximately 850K CpGs, is likely to be used in many future methylation data collection efforts. Since genotype data and corresponding EPIC array data for the same individuals were not publicly available at the time of this study, we were not able to compile a reference list of CpGs for the EPIC array. However, inspection of the probes available in each array reveals that only 32,425 of the probes in the 450K array were not included in the EPIC array. We further found that 94% of the CpGs in the 450K-based reference list we constructed (4,616 CpGs out of 4,913) were included in the EPIC array. Therefore, our suggested 450K-based reference list is expected to perform similarly on data generated from the EPIC array. In order to test that, we repeated all of the experiments we performed so far, only this time we removed from the data the set of 32,425 sites that were not included in the EPIC array. The results, summarized in Supplementary Table S1, show that removing these sites leads to only a marginal decrease in the *R*^2^ values. Clearly, as more EPIC array data will become available, EPIC-based reference list of CpGs is expected to further improve the performance of EPISTRUCTURE.

## Discussion

We demonstrated that 450K DNA methylation data can capture population structure in admixed populations. In particular, we observed that in the presence of a relatively strong population structure (GALA II) the dominant genome-wide signal of ancestry information could be revealed once appropriately correcting for tissue heterogeneity. In contrast, we observed that in the presence of weaker population structure in the data (CHAMACOS) the genome-wide signal of ancestry methylation is only moderately reflected by the dominant axes of variance in the data after accounting for tissue heterogeneity.

Using KORA, a large data set for which both methylation levels and genotypes were available, we generated a reference list of genetically-informative CpGs and successfully used it to estimate ancestry information in new data sets by applying PCA on the reference sites. Polymorphic CpGs that were found to be highly correlated with genetics were also include in the reference list. Although these CpGs are generally teated as artifacts, they represent true genetic signal and therefore were used in order to further increase the signal captured by EPISTRUCTURE. As we showed, by taking this approach, EPISTRUCTURE was able to effectively isolate and capture ancestry information in methylation data.

While we observed strong correlations between the EPISTRUCTURE PCs and the genotype-based population structure estimates of the GALA II individuals, only moderate correlations were found in the CHAMACOS data set (though substantially better than unsupervised approaches, in which only negligible correlations to the true ancestry were found). These results can be explained by the fact that only 106 AIMs were available for us in the CHAMACOS for capturing ancestry information, as opposed with the dense genotype array information used in the GALA II analysis. Therefore, it is likely that our inference of population structure by methylation data is in fact more accurate than reflected in the experiments conducted on the CHAMACOS samples.

The reference-list of CpGs was generated using methylation states and genotypes collected from European individuals, therefore it may not be optimized for capturing ancestry information in non-European populations. However, since we successfully used this list for the inference of ancestry in the Latino GALA II and CHAMACOS individuals, we expect it to prove useful for some other non European populations as well. We further note that when constructing the linear models for each CpG from its cis-SNPs in the whole-blood KORA data, we decided not to account for tissue heterogeneity. To the best of our knowledge, there is currently no evidence for dramatic genome-wide effects of genotypes on the cell type composition. Therefore, in the vast majority of CpGs, the cell type composition is expected to be orthogonal to the genetic signal they contain. As a result, accounting for tissue heterogeneity in this case is more likely to reduce the accuracy of the model due to inaccuracies of the cell type composition estimates rather than to bias the selection of CpGs into the reference-list.

## Conclusions

Genome-wide DNA methylation levels reflect ancestry information, which can be effectively isolated using CpGs that are highly correlated with their cis-SNPs. In line with previous works showing many associations of methylation with genetic variation and ancestry, our results further emphasize the importance of accounting for ancestry information in methylation studies of diverse populations. In the absence of genotype data we suggest that our proposed method, EPISTRUCTURE, can be used in EWAS for accounting for population structure.

## Methods

### Data and quality control

The longitudinal KORA study (Cooperative health research in the Region of Augsburg) consists of independent population-based subjects from the general population living in the region of Augsburg, southern Germany [25]. Whole blood samples of the KORA F4 study were used (*n* = 1, 799) as described elsewhere [39]. Briefly, DNA methylation levels were collected using the Infinium HumanMethylation450K BeadChip array (Illumina). Beta Mixture Quantile (BMIQ) [40] normalization was applied to the methylation levels using the R package wateRmelon, version 1.0.3 [41]. In total 431,360 probes were available for the analysis. As described elsewhere [42], genotyping was performed with the Affymetrix 6.0 SNP Array (534,174 SNP markers after quality control), with further imputation using HapMap2 as a reference panel. A total of 657,103 probes remained for the analysis.

We used whole-genome DNA methylation levels and genotyping data from the Genes-environments & Admixture in Latino Americans (GALA II) data set, a pediatric Latino population study. Details of genotyping data including quality control procedures for single nucleotide polymorphisms (SNPs) and individuals have been described elsewhere [38]. Briefly, participants were genotyped at 818,154 SNPs on the Axiom Genome-Wide LAT 1, World Array 4 (Affymetrix, Santa Clara, CA) [43]. Non-autosomal SNPs and SNPs with missing data (> 0.05) and/or failing platform-specific SNP quality criteria (*n* = 63, 328) were excluded as well as SNPs not in Hardy-Weinberg equilibrium (*n* = 1, 845; *p* < 10^−6^) within their respective populations (Puerto Rican, Mexican, and other Latino). Study participants were filtered based on 0.95 call rates and sex discrepancies, identity by descent and standard Affymetrix Axiom metrics. Finally, SNPs with low MAF (< 0.05; *n* = 334, 975) were excluded. The total number of SNPs passing QC was 411,787. The data are available in dbGaP (accession ID phs000920.v1.p1).

Whole-blood methylation data for a subset of the GALA II participants (*n* = 573) are publicly available in the Gene Expression Omnibus (GEO) database (accession number GSE77716) and have been described elsewhere [13,23]. Briefly, methylation levels were measured using the Infinium HumanMethylation450K BeadChip array and raw methylation data were processed using the R minfi package [44] and assessed for basic quality control metrics, including determination of poorly performing probes with insignificant detection P-values above background control probes and exclusion of probes on X and Y chromosomes. Finally, beta-normalized values of the the data were SWAN normalized [45], corrected for batch using COMBAT [46] and adjusted for age, gender and chip assignment information using linear regression. The number of participants with both methylation and genotyping data was 525. We further excluded 46 individuals collected in a separate batch since they were all Puerto-Ricans. A total of 479 individuals and 473,838 probes remained for the analysis.

In order to further evaluate and validate the performance of EPISTRUCTURE we used data from the CHAMACOS longitudinal birth cohort study [34]. For this analysis, we had a subset of subjects that had Infinium HumanMethylation450K BeadChip array data available at 9 years of age. Briefly, samples were retained only if 95% of the sites assayed had insignificant detection P-value and samples demonstrating extremes level in the first two PCs of the data were removed. Probes where 95% of the samples had insignificant detection P-value (> 0.01; *n* = 460) as well as cross-reactive probes (*n* = 29, 233) identified by Chen et al. [24] were dropped. A total of 227 samples and 455,590 probes remained for the analysis. Color channel bias, batch effects and difference in Infinium chemistry were minimized by application of All Sample Mean Normalization (ASMN) algorithm [47], followed by BMIQ normalization [40]. The data were adjusted for gender and technical batch information using linear regression.

In line with a previous study showing that a panel of small size is sufficient to approximate genetic ancestry in Latino populations well [48], 106 SNPs were collected and used as AIMs for estimating genetic ancestry of the CHAMACOS individuals [36]. The panel of AIMs was selected according to previously reported studies of Latino populations [12,36,49,50]. Briefly, only SNPs with large differences in allele frequencies between ancestries were selected.

### 450K Human Methylation array

This state of the art methodology allows for examination of > 450, 000 CpG sites across the genome, representing 99% of RefSeq genes. Sites include promoters, gene bodies, and 96% of UCSC database CpG islands (dense concentrations of CpGs), many of which are known to be associated with transcriptional control [51–56]. This platform has been especially amenable to population studies because of its relative cost effectiveness and low sample requirements. Several studies have identified CpG sites differentially methylated by environmental exposures [57, 58] (e.g. arsenic and tobacco smoke) and health outcomes including obesity [59], rheumatoid arthritis [60], and Crohns disease [61] demonstrating its utility in environmental and molecular epidemiology studies. The relative methylation (beta-normalized values) for each CpG site is calculated as the ratio of methylated-probe signal to total (methylated + unmethylated) fluorescent signal intensity. The Infinium pipeline is streamlined with excellent reproducibility [62].

### Model and algorithm

Previous studies reported a large number of correlations between DNA methylation and genetics, mainly cis-correlations between CpGs and nearby SNPs [9–12]. We therefore assume that cis-located SNPs can capture the genetic variability accounting for the methylation levels of a given CpGs. For a given CpG *m* we assume the following linear model:

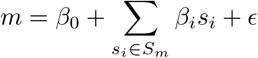

where *S*_*m*_ is a group of *w* SNPs, cis-located with respect to *m*, *β*_i_ values are their corresponding effects on the methylation levels and ∈ represents an error term, assumed to be independent between different samples.

Given reference data of methylation levels and genotypes for the same individuals, we fit the above linear model for each CpG. We regard the CpGs for which the model fits well as linear combinations of SNPs. We define the set of these genetically-informative CpGs as the reference list. Given methylation data for new individuals, we can estimate the population structure in the data by applying a standard PCA on the sites in the reference list. The first several PCs of PCA are well-known to efficiently capture ancestry information when applied to genotypes data [4], therefore applying PCA on CpGs that are linear combinations of SNPs is expected to capture population structure as well (see Supplementary note for more details).

Given reference methylation and genotypes data, our suggested algorithm can be summarized as follows:

1. For each CpG *m*, fit a linear model using *w* SNPs that are closest to *m*.
2. Define a reference list *G* of all the CpGs for which the linear model fits well. Evaluate model fit based on cross-validated squared linear correlation.
3. Given a new methylation data set, apply PCA on the sites defined by *G* and consider the first *k* principal components as the estimate of the population structure.

Note that creating a reference list, described in the first two steps of the algorithm, needs to be performed only once. Population structure can be then inferred in future data sets using this list of genetically-informative CpGs. In practice, an appropriate *w* may be relatively large (i.e. large number of predictors), while the sample size is typically limited. We therefore apply a regularized regression with ℓ_1_ penalty, also known as LASSO regression [63]. For the same reason, we define a parameter *p* to limit the maximal number of predictors in each model. Furthermore, in order to avoid over-fitting of the model, we perform a k-fold cross-validation procedure for each CpG. The score of a CpG is defined as the median squared linear correlation of its predicted values with the real values across the *k* folds. Finally, a reference list of the CpGs is defined as the set of sites with highest scores.

In principle, one could use the same approach taken here in order to create and use a reference list of CpGs which explain SNPs well rather than CpGs which are captured well by SNPs. However, modeling methylation levels as a function of SNPs is more natural with respect to the causality relations assumed between SNPs and methylation. Moreover, many methylation sites are known to be affected by several factors (e.g. age [28], gender [29] and smoking [64]), and therefore considering a group of methylation sites explaining a given SNP may introduce into the data more, potentially unknown, variance in addition to genetic variance. This potential problem is expected to be less severe in the opposite direction of modeling methylation using SNPs. In this case, methylation sites that are well explained by genetics are less likely to be highly explained by more factors.

### Compiling a reference list from the KORA cohort

The reference list of genetically-informative CpGs was created using the KORA cohort for which whole-blood methylation data as well as genotype data were available for 1,799 European individuals. Following the algorithm described above, a score was computed for each CpG using k-fold cross-validation with *k* = 10 and using the parameters *w* = 50 and *p* = 10 (Additional file 2). A reference list was then compiled from CpGs with median correlation of *R*^2^ > 0.5 in the cross-validated prediction procedure, resulting in a total of 4,913 CpGs (Additional file 3), out of which 2,436 are polymorphic CpGs and additional 801 CpGs have at least on common SNP in their probe outside the CpG target. The number of these reference CpGs available in the GALA II data set and in the CHAMACOS data set were 4,912 and 4,450, respectively. Removing probes with polymorphic CpGs results in 2,476 and 2,229 CpGs, and further removing probes with common SNPs results in 1,676 and 1,554 CpGs for GALA II and CHAMACOS, respectively. Unless stated otherwise, polymorphic CpGs were not excluded from the reference of informative CpGs in the executions of EPISTRUCTURE, therefore highly informative polymorphic CpGs (*R*^2^ > 0.5) were also included in the reference list. In most cases, polymorphic CpGs are excluded as a preprocessing step in epigenetic studies, however, here we leverages the true genetic signal underlying in these probes for capturing the ancestry information better.

### Detecting 450K probes containing SNPs

Probes with a SNP in their CpG target (polymorphic CpGs) were shown to be biased with underlying genetic polymorphisms rather than capture methylation signals solely [24]. The authors reported a list of 70,889 such polymorphic CpGs in the 450k DNA methylation array, as well as a list of common SNPs residing in probes of the 450K array outside the CpG target (MAF > 0.01 according to at least one of the major continental groups in the 1000 Genome database [65]). The total number of probes containing SNPs reported is 167,738.

### Estimating ancestry information

Proportions of European, Native-American and African ancestries were estimated for each individual in both the GALA II and the CHAMACOS cohorts using the software ADMIXTURE [5] and the default reference data provided by the software. For the GALA II individuals we used the 411,787 SNPs remained after QC as an input, and for the CHAMACOS individuals we used the 106 available AIMs. The genotype based PCs were computed by applying PCA on the standardized values of the available genotypes in each data set. For the CHAMACOS data set, prior to computing PCA, we excluded sites with more than 5% missing values and completed the remaining missing values by assigning the mean. This resulted in a total of 99 SNPs.

### Adjusting methylation levels for tissue heterogeneity

Methylation levels of the GALA II and CHAMACOS data sets were adjusted for cell type composition using ReFACTor, a reference-free method for the correction of cell type heterogeneity in EWAS [23]. Each data set was adjusted for cell composition by regressing out the first six ReFACTor components, resulting in adjusted beta values. ReFACTor was executed using the default parameters and *K* = 6. For one of the experiments in the GALA II data we used an alternative approach for cell type composition correction. Similarly, as with the ReFACTor components, we generated beta adjusted values, only this time we used reference-based cell proportion estimates of main leukocyte cell types. Specifically, we obtained cell proportion estimates of six cell types (granulocytes, monocytes, B cells, NK cells, CD8T and CD4T cells) using the default implementation available in the minfi package [44], defined and assembled for the 450K array [66] based on the approach suggested by Houseman et al [32] and a 450K reference data set [67].

### Feature selection based on proximity to SNPs

For evaluating our suggested method, we calculated alternative methylation-based PCs after applying a feature selection that was previously suggested as a method for capturing population structure [21]. Following the authors’ recommendation, we considered a list of the CpGs residing within 50 bp from SNPs, as provided by the authors.

## Acknowledgments

This research was partially supported by the Edmond J. Safra Center for Bioinformatics at Tel-Aviv University. E.H., E.R. L.S. and R.S. were supported in part by the Israel Science Foundation (Grant 1425/13), E.H., E.R., and R.S. by NSF grant 1331176 and United States Israel Binational Science Foundation grant 2012304. E.R. and L.S. were supported by Len Blavatnik and the Blavatnik Research Foundation. C.E., S.H., D.H., J.G., S.O., E.B. were supported by the Sandler Family Foundation, the American Asthma Foundation, Hind Distinguished Professorships, and by the following research grants: 1P60 MD006902, 1R01HL117004-01, R21ES24844-01, U54MD009523, TRDRP 24RT-0025. N.Z. was supported in part by an NIH career development award from the NHLBI (K25HL121295). KORA was initiated and financed by the Helmholtz Zentrum MnchenGerman Research Center for Environmental Health, Neuherberg, Germany and supported by grants from the German Federal Ministry of Education and Research (BMBF), the Federal Ministry of Health (Berlin, Germany), the Ministry of Innovation, Science, Research and Technology of the state North RhineWestphalia (Dsseldorf, Germany), and the Munich Center of Health Sciences (MC Health) as part of LMUinnovativ. We thank all members of field staffs who were involved in the planning and conduct of the MONICA/KORA Augsburg studies. The funders had no role in study design, data collection, and analysis, decision to publish, or preparation of the manuscript. The research leading to these results has received funding from the European Union Seventh Framework Programme (FP7/20072013) under grant agreements n 261433 (BioSHaRe) and n 603288 (sysVASC) and was supported by the European Unions Seventh Framework Programme (FP7HealthF52012) under grant agreement no. 305280 (MIMOmics), the HelmholtzRussia Joint Research Group (HRJRG) 310, the German-Israeli Foundation for Scientific Research and Development (Grant 109433.2/2010) and the German Center of Diabetes Research (DZD). The CHAMACOS study was supported by the NIH grants P01 ES009605, RO1 ES012503, R01 ES 023067 and R01 ES021369, and EPA grants RD 82670901 and RD 83451301. The contents of this manuscript are solely the responsibility of the authors and do not necessarily represent the official views of the NIEHS and the EPA.

